# Unexpected Loss of Sensitivity to the nAChR Antagonist Activity of Mecamylamine and DHβE in Nicotine-Tolerant C57BL/6J Mice

**DOI:** 10.1101/482075

**Authors:** Fernando B. de Moura, Lance R. McMahon

## Abstract

There has always been interest in developing nAChR antagonists as smoking cessation aids, to add to nAChR agonists (e.g., nicotine replacement) already used for that indication. Previous studies have demonstrated that daily nicotine treatment confers tolerance to some of the effects of nicotine, as well as cross-tolerance to other nAChR agonists. The current study assessed the extent to which antagonism of nicotine varies as a function of daily nicotine treatment. The rate-decreasing and hypothermic effects of nicotine, as well as antagonism of those effects, were examined in C57BL/6J mice before, during treatment with, and after discontinuation of three daily injections of 1.78 mg/kg nicotine. The nonselective nAChR antagonist mecamylamine and the β2 nAChR antagonist DHβE were studied in combination with nicotine. The ED_50_ values of nicotine to produce rate-decreasing and hypothermic effects were, respectively, 0.44 and 0.82 mg/kg prior, 1.6 and 3.2 mg/kg during, and 0.74 and 1.1 mg/kg after discontinuation of daily nicotine treatment. Prior to daily nicotine treatment, mecamylamine decreased response rate and rectal temperature; however, during daily nicotine, mecamylamine (up to 5.6 mg/kg) only decreased rectal temperature. DHβE (up to 5.6 mg/kg) when studied prior to daily nicotine decreased rectal temperature, but that decrease was abolished during chronic nicotine treatment. Mecamylamine and DHβE antagonized the rate-decreasing and hypothermic effects of nicotine before and after daily nicotine; however, during daily nicotine, mecamylamine and DHβE antagonized only the hypothermic effects of nicotine. The differential antagonism of rate-decreasing and hypothermic effects implicates differential involvement of nAChR subtypes. The decreased capacity of mecamylamine and DHβE to antagonize nicotine during chronic nicotine treatment may indicate that their effectiveness as smoking cessations might vary as a function of nicotine tolerance and dependence.

## Introduction

Tobacco use is the leading cause of preventable death (World Health Organization 2011). Current smoking cessation aids improve abstinence outcomes, although there is marked room for improvement inasmuch as 75% of individuals relapse to the use of tobacco products within one year of initiating pharmacotherapy (Hays et al. 2008). There is a growing interest in new strategies to combat the tobacco epidemic, including the use of both drug and behavioral interventions (Brunzell et al. 2014; Mooney et al. 2016; Vogeler et al. 2016; Windle et al. 2016). A potential strategy previously considered but not yet formally approved is the use of nicotinic acetylcholine receptor (nAChR) antagonists, which have been demonstrated to attenuate the reinforcing effects of nicotine as well as nicotine-induced increases in mesolimbic dopamine (Corrigall et al. 1992; Nisell et al. 1994; Crooks et al. 2014). One such antagonist is the nonselective, noncompetitive nAChR ligand mecamylamine (Lancaster and Stead 2000; Frishman et al. 2006). The value of mecamylamine as a smoking cessation aid was underscored by reports that mecamylamine did not induce withdrawal in human tobacco users (Eissenberg et al. 1996), which if it had would have created concerns regarding compliance. However, mecamylamine, and other antagonists, have not been widely used as smoking cessation aids. Studies have suggested that mecamylamine, when used alone, is only effective in approximately 15% of participants in clinical studies (Rose et al. 1994; Lancaster and Stead 2000).

One issue to be considered when developing any pharmacotherapy for drug abuse is how chronic treatment with an abused drug alters the receptor systems mediating the effects of not only the abused drug, but also a pharmacotherapy that may act through the same receptor systems. In the same way daily tobacco use confers tolerance to many of the effects of nicotine and other nAChR agonists (Rosecrans et al. 1989; Cunningham and McMahon 2011; Rodriguez et al. 2014; de Moura and McMahon 2016), the effects of nAChR antagonists might also be impacted. Repeated exposure to nicotine differentially impacts the expression levels of various nAChR subtypes within the CNS (Olale et al. 1997; Fenster et al. 1999; Buisson and Bertrand 2001; Nashmi et al. 2007). Whether neuroadaptations that occur as a result of nicotine exposure can impact the effects of nAChR antagonists, particuraly their nicotine antagonist activity, has not been fully characterized.

The nonselective nAChR antagonist mecamylamine (Banerjee et al. 1990) and the selective β2 nAChR antagonist dihydro-β-erythroidine (DHβE) (Papke et al. 2008) have been studied under conditions of chronic nicotine exposure to examine precipitated withdrawal (Damaj et al. 2003; Vann et al. 2006). However, to our knowledge, there is no published study that has examined the extent to which antagonism of nicotine varies as a function of chronic nicotine treatment. This study identified whether exposure to nicotine under conditions that produce tolerance, but are unlikely to be sensitive to antagonist-induced precipitated withdrawal (Damaj et al. 2003; de Moura and McMahon 2016), disrupts the ability of the nAChR antagonists mecamylamine and DHβE to block the *in vivo* effects of nicotine. Mecamylamine was selected as a test drug because it is a nonselective nAChR antagonist which has been studied in human clinical studies as a potential smoking cessation aid, albeit with limited positive results (Lancaster and Stead 2000; Frishman et al. 2006; Crooks et al. 2014). DHβE was selected as a test drug because the β2 subunit has been demonstrated to mediate the reinforcing effects of nicotine and nicotine-induced dopamine release (Picciotto et al. 1998; Salminen et al. 2007; Crooks et al. 2014). Simple schedule-controlled responding and hypothermia were selected for study because they have been previously used to examine the pharmacology of nicotine, and both are sensitive to the development nicotine tolerance (Rosecrans et al. 1989; Cunningham and McMahon 2011; Rodriguez et al. 2014; de Moura and McMahon 2016).

## Methods

### Subjects

Male C57BL/6J mice were purchased at 8 weeks of age (n=8; Jackson Laboratories, Bar Harbor, Maine, USA) and were housed individually in a temperature, controlled room (23°C), under a 14/10h light/dark cycle. Mice were food restricted to 85% of their free-feeding body weight, while water was available ad libitum. Food (Dustless Precision Pellets Grain-Based Rodent Diet; Bio-Serv, Frenchtown, New Jersey, USA) was available immediately following experimental sessions. Experimental conditions were in accordance with those set forth by the National Institute of Health’s Guide for the Care and Use of Laboratory Animals (Institute of Laboratory Animal Research, 2011). The Institutional Animal Care and Use Committee of University of Texas Health San Antonio approved these experiments.

### Apparatus

Operant conditioning chambers (ENV-307A-CT; Med Associates, St. Albans, Vermont, USA) were kept in sound-attenuating and ventilated boxes. Each chamber contained a light and a recessed 2.2 cm-diameter hole on one wall through which reinforcers could be presented. On the opposite wall were three identical holes arranged horizontally, spaced 5.5 cm apart located. The centers of each hole were 1.6 cm from the floor of the chamber. During experimental sessions, when the middle hole was illuminated, a mouse could disrupt a photobeam by a nose-poke, resulting in the presentation of a 0.01 ml of 50% v/v unsweetened condensed milk/water through the hole on the opposite wall. A computer was connected to the operant conditioning chambers through an interface (MED-SYS-8; Med Associates), and experimental sessions were controlled, and responses were recorded, using Med-PC software (Med Associates). Rectal temperature was recorded using a rectal probe designed for mice attached to a digital thermometer (BAT7001H; Physitemp, Clifton, New Jersey, USA). The probe length was 2 cm. The probe diameter was 0.7112 mm, except for the tip, which was 1.651 mm (RET-2-ISO; Physitemp, Clifton, New Jersey, USA). The probe was inserted 2 cm into the rectum.

### Experimental Procedure

After 7 days of habituation to the housing room, experimental sessions were conducted 7 days per week during the light cycle. Mice were shaped to respond in the middle hole during 60-min sessions. One response resulted in 10-s access to milk. When milk was presented, a house light was illuminated and the light in the middle hole was extinguished. Responses during presentation of the milk reinforcer had no programmed consequence. When 6 of 8 mice obtained 100 reinforcers for three consecutive sessions, the session length was shortened to 25 min. Sessions were divided into a 10-min pretreatment where the mouse was in the operant chamber but no lights were illuminated and responses had no programmed consequence, followed by 15 min of milk availability in the presence of light. The response requirement was systematically increased to a fixed ratio 20 (FR20). Mice received saline immediately prior to these sessions.

When the rate of responding for 6 mice was within ±20% of baseline, defined as the running average of the previous 5 sessions, for 5 consecutive, or 6 of 7 sessions, drugs were administered at the beginning of sessions. One dose or a combination of doses was administered per drug test, and all mice received drugs on the same day. After drug tests, saline was administered during intervening sessions. Before the next drug test, the response rate of 6 mice was required to be within ±20% of baseline for at least 2 out of 3 sessions. Immediately before and after every experimental session, rectal temperature was measured. Dose-effect functions for rate-decreasing and hypothermic effects were generated in the following order: nicotine, mecamylamine, DHβE, and nicotine in the presence of DHβE. Doses were administered in a non-systematic order among the mice so that no more than 2 mice received the same dose or dose combination for that particular test. The dose-effect function for nicotine was re-determined twice: once before studies with mecamylamine and DHβE, and once immediately afterwards.

The timeline in Figure 1 shows the daily sequence (i.e., injections, temperature measurement, and operant session) during the course of daily nicotine exposure. After the second nicotine dose-effect determination, nicotine was administered daily in three separate doses (1.78 mg/kg each dose) spaced 90 min apart. Rectal temperature was measured immediately before and 30 min after each dose. For the first 10 days of daily nicotine treatment, operant sessions were conducted following the third nicotine dose (Figure 1a). However, one of the mice did not respond. Therefore, starting on day 11, the 25-min operant session was conducted 1 h after the second nicotine dose, 30 min prior to the third nicotine dose (Figure 1b). Saline or drug were administered immediately before the operant sessions. Sessions were conducted daily until the rate of responding for 6 mice did not vary by ±20% of baseline for 5 consecutive, or 6 of 7, sessions. Thereafter, dose-effect functions were determined for nicotine, mecamylamine, and DHβE. Moreover, the nicotine dose-effect function was determined in the presence of mecamylamine or DHβE. The order of tests with an antagonist alone or nicotine in combination with an antagonist was non-systematic among mice, and dose-effect data were collected between days 16 and 100 of chronic nicotine exposure. On days 43-59 of chronic nicotine exposure, mice received three doses of 1.78 mg/kg nicotine daily, but experimental sessions were not conducted. After daily nicotine exposure was discontinued, mice were treated with saline instead of nicotine three times daily. Sensitivity to 1 mg/kg nicotine was examined every 8 sessions. Once sensitivity to the effects of 1 mg/kg nicotine was stable (i.e., not significantly different) for 3 consecutive tests, dose-effects functions for nicotine, nicotine in the presence of mecamylamine, and nicotine in the presence of DHβE were re-determined.

**Figure 1:**
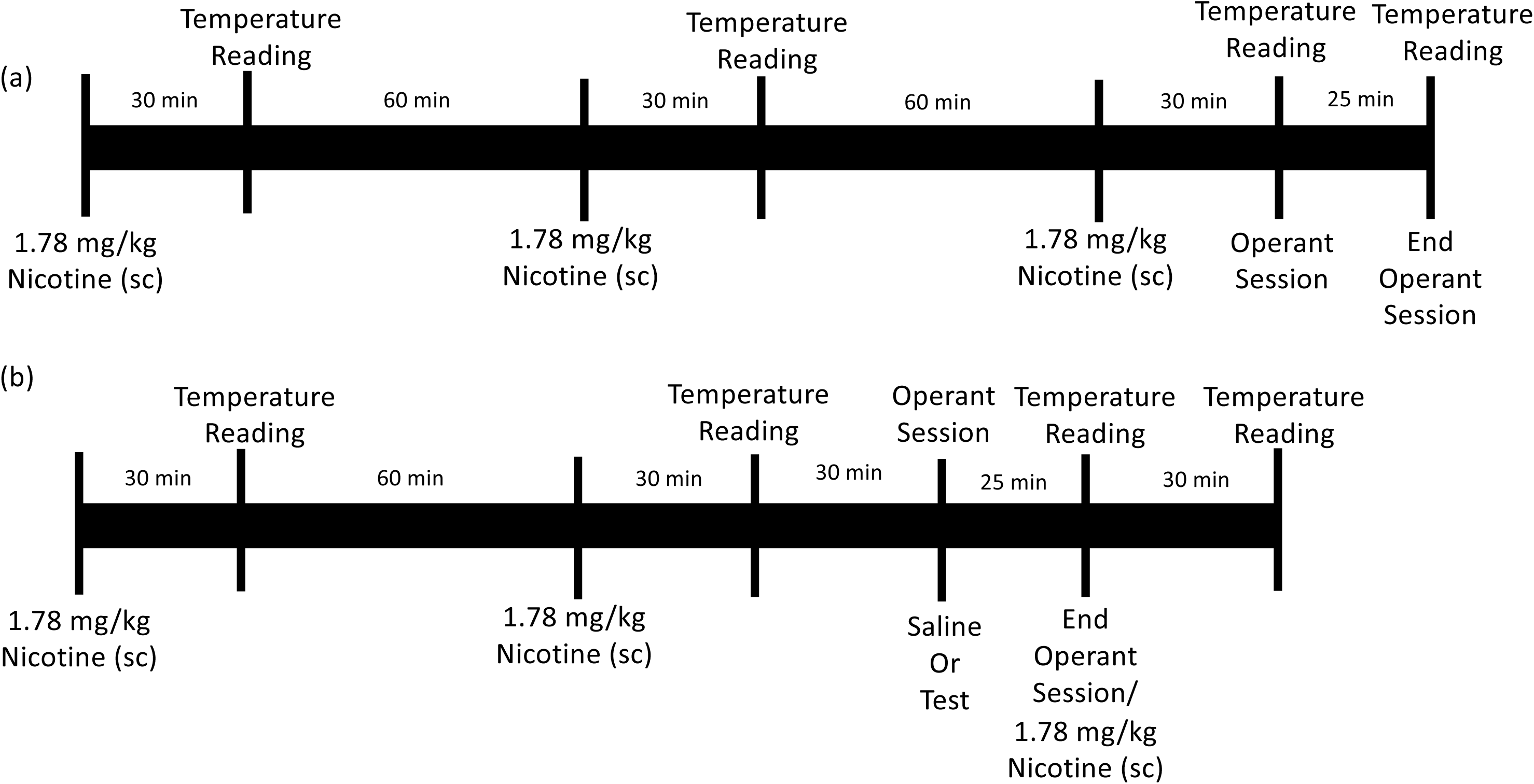
Schematic of experimental procedure during (a) days 1-10 and (b) days 11-100 of daily nicotine exposure. On days 1-10, 1.78 mg/kg nicotine was administered subcutaneously every 90 min, with a rectal temperature measurement 30 min after nicotine administration. Immediately following the third nicotine dose, the mice were placed in the operant chamber for a 25-min experimental session; upon termination of the session, rectal temperature was measured again. On days 11-100, 1.78 mg/kg nicotine was administered every 90 min, and rectal temperature was measured 30 min after nicotine administration. However, operant sessions were conducted 60 min after the second nicotine dose, 30 min before the third nicotine dose. Operant sessions were conducted immediately following saline, or a dose of nicotine alone, or in combination with an antagonist. Only one test session was conducted per day. Rectal temperature was measured immediately upon termination of the operant session, and was followed by the third injection of nicotine only if saline preceded the experimental session.

### Drugs

The drugs were nicotine hydrogen tartrate salt (Sigma-Aldrich, St. Louis, Missouri, USA), mecamylamine hydrochloride (Waterstone Technology, LLC, Carmel, Indiana, USA), and dihydro-β-erythroidine hydrobromide (DHβE; Tocris Biosciences, Bristol, UK). Nicotine and DHβE were administered subcutaneously, while mecamylamine was administered intraperitoneally. DHβE and mecamylamine were administered 5 min prior to operant sessions. All drugs were administered in a volume of saline equal to 10 ml/kg; nicotine solutions were adjusted to pH of 7. Drug doses are expressed as the combined weight of base and salt, except for nicotine, which is expressed as the base weight.

### Data Analyses

Data were plotted as mean ± standard error of the mean and analyzed as responses per s and change in °C, except for the analysis of DHβE in combination with nicotine during chronic nicotine treatment. For that analysis, data were expressed as a percentage of the individual saline-control response rate and as a change in °C from the individual saline-control rectal temperature. Saline control was defined as the running average of the 5 previous sessions during which saline was administered. Saline controls determined before, during, and after discontinuation of chronic nicotine treatment were compared with a one-way repeated measures ANOVA followed by a Dunnett’s test (*p*<0.05). Response rate following the daily nicotine dose (1.78 mg/kg) was examined with a one-way repeated measures ANOVA, with consecutive days of treatment as the main factor, followed by a Dunnett’s test. A two-way repeated measures ANOVA followed by Tukey’s multiple comparison tests was used to analyze hypothermic effects, with daily nicotine dose (i.e., first, second, and third) as one factor and consecutive days of treatment was a second factor.

Straight lines were fitted to nicotine dose-effect data using linear regression (GraphPad Prism version 6.0 for Windows, GraphPad Software, San Diego, California, USA). The linear portion of the function was determined per mouse. For response rate, the function included the largest dose producing no more than a 20% decrease up to and including the smallest dose producing greater than an 80% decrease. For rectal temperature, the function included the largest dose resulting in less than a 1°C change up to and including the smallest dose producing greater than a 5°C change. The ED_50_ values and potency ratios were calculated from individual nicotine dose-effect functions using a common best-fitting slope (Tallarida, 2000). The ED_50_ values were significantly different when the 95% confidence limits calculated from the individual potency ratios did not include 1. Because mecamylamine and DHβE alone, up to the largest doses studied, were less effective than nicotine in decreasing response rate and rectal temperature, dose-effect data were analyzed separately for each antagonist with one-way repeated measures ANOVAs followed by Dunnett’s tests.

One-way repeated measures ANOVAs followed by a Dunnett’s test was used to compare the individual nicotine ED_50_ values determined before, during, and after discontinuation of treatment. Student’s *t*-test was used to compare potency ratios (i.e., magnitude of antagonism by DHβE) before and after discontinuation of chronic nicotine treatment. During chronic treatment, only the smallest doses of nicotine (0.56 and 1 mg/kg) appeared to be altered by mecamylamine or DHβE relative to the nicotine control; these dose-effect data were analyzed separately per antagonist with two-way repeated measures ANOVAs, with nicotine dose being one factor and saline versus antagonist being a second factor. Dunnett’s tests were used to compare the effects of nicotine to saline, or nicotine in combination with an antagonist to the effects of the antagonist alone. Sidak’s tests were used to compare the effects of each dose of nicotine in the presence versus the absence of antagonist. A Student’s t-test was used to compare the effects of saline to a dose of antagonist.

## Results

The effects of saline administered at the beginning of operant sessions were compared at each phase of the study including before, during, and after discontinuation of daily nicotine treatment. Once stable responding was achieved, the respective averages (± SEM) in responses per s were 0.68±0.10, 0.51±0.07, and 0.59±0.11. These response rates were not significantly different from one another (F_2,12_=3.4, p=0.09). The respective rectal temperatures measured after these sessions were 37.1±0.07, 36.7±0.04, and 36.5±0.11 °C. These temperatures were significantly different from one another (F_2,12_=30.7, p=0.0001); rectal temperature before daily nicotine exposure was significantly higher than rectal temperature during and after daily nicotine exposure.

The nicotine dose-effect functions, one determined before tests with DHβE and the second just before chronic nicotine treatment, were not significantly different from each other for response rate (F_2,42_=2.8, p=0.071) and hypothermia (F_2,42_=1.98, p=0.15). Therefore, the nicotine dose-effect functions were averaged to produce a single control for further analyses. Nicotine dose-dependently decreased response rate and rectal temperature (Figure 2); after saline pretreatment, 1.78 mg/kg nicotine decreased response rate to 0.0004 responses per s and decreased rectal temperature to 31.1°C (Figure 2, circles). The ED_50_ values (95% confidence limits) of nicotine to decrease response rate and rectal temperature were 0.44 (0.32-0.61) and 0.82 (0.72-0.94) mg/kg, respectively. DHβE (3.2 mg/kg) alone resulted in an average response rate and rectal temperature of 0.80±0.16 responses per s and 37.2±0.07°C, which were not significantly different from saline (t_7_=1.2, p=0.25 and t_7_=0.08, p=0.94, respectively) (Figure 2, leftmost squares). DHβE shifted the nicotine dose-effect functions for decreasing response rate and rectal temperature 2.9 (2.2-3.9) fold and 3.2 (2.7-3.8) fold to the right, respectively (Figure 2, squares).

**Figure 2:**
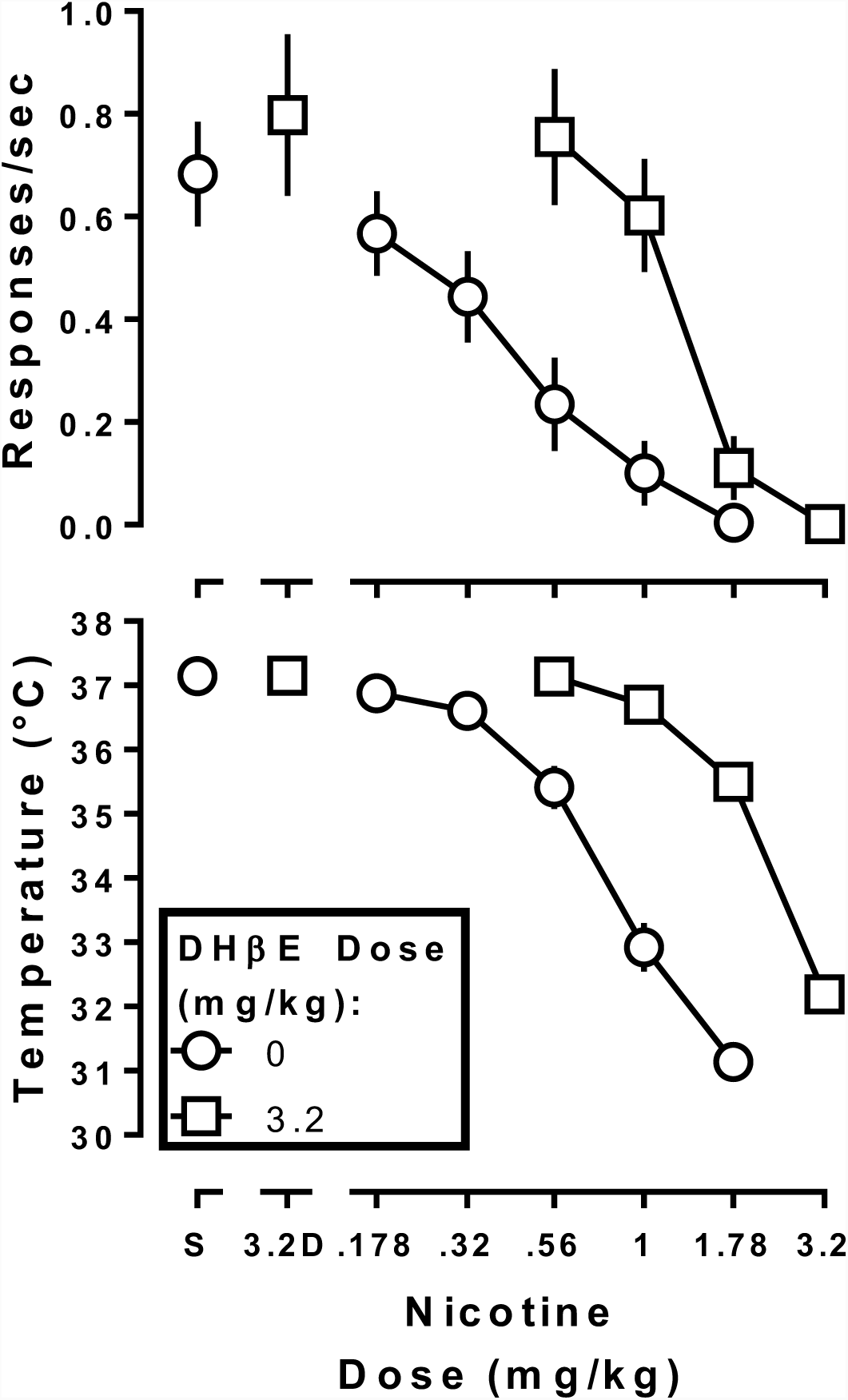
Rate of responding (top panel) and rectal temperature (bottom panel) following nicotine alone and in combination with 3.2 mg/kg DHβE (squares) prior to daily nicotine exposure. Ordinate: response rate expressed responses/s (top panel), and rectal temperature expressed as °C (bottom panel). Abscissa: nicotine dose in mg of the free base per kg body weight.

On day 1 of daily nicotine treatment, response rate determined 10 min after the third dose (1.78 mg/kg) of nicotine was 0.44 responses per s, a decrease of 37% relative to the saline control determined before daily nicotine treatment (Figure 3, top). Responding after the last daily nicotine injection systematically decreased on subsequent days, was lowest on day 4, and remained low (F_6,11_=14, p=0.0016) until the timing of sessions with nicotine injections was changed so that sessions were conducted 1 h after the second daily dose of nicotine beginning on day 11 (Figure 1, compare top and bottom timelines). The first dose of nicotine on the first day of treatment decreased rectal temperature to 30°C. The second and third doses of nicotine also decreased rectal temperature (i.e., to 32 and 33°C, respectively), although the hypothermic effects of nicotine after the second and third nicotine daily doses were less than the first dose (Figure 3, bottom, day 1). Tolerance developed to the hypothermic effects produced by each dose of nicotine across days of repeated, daily nicotine treatment (F_21,126_=54, p<0.0001). Rate of responding was stable according to the established criterion on day 16 (Figure 3, top). During daily nicotine, the nicotine dose-effect functions were significantly shifted to the right (Figure 4, black circles); the ED_50_ values for nicotine to produce rate-decreasing and hypothermic effects were 1.6 (1.4-1.9) and 3.2 (2.7-3.8) mg/kg, respectively. Daily nicotine exposure produced a 3.6 (2.6-5.0) and 4.8 (3.9-5.9) fold rightward shift in the nicotine dose-effect functions for rate-decreasing and hypothermic effects, respectively.

**Figure 3:**
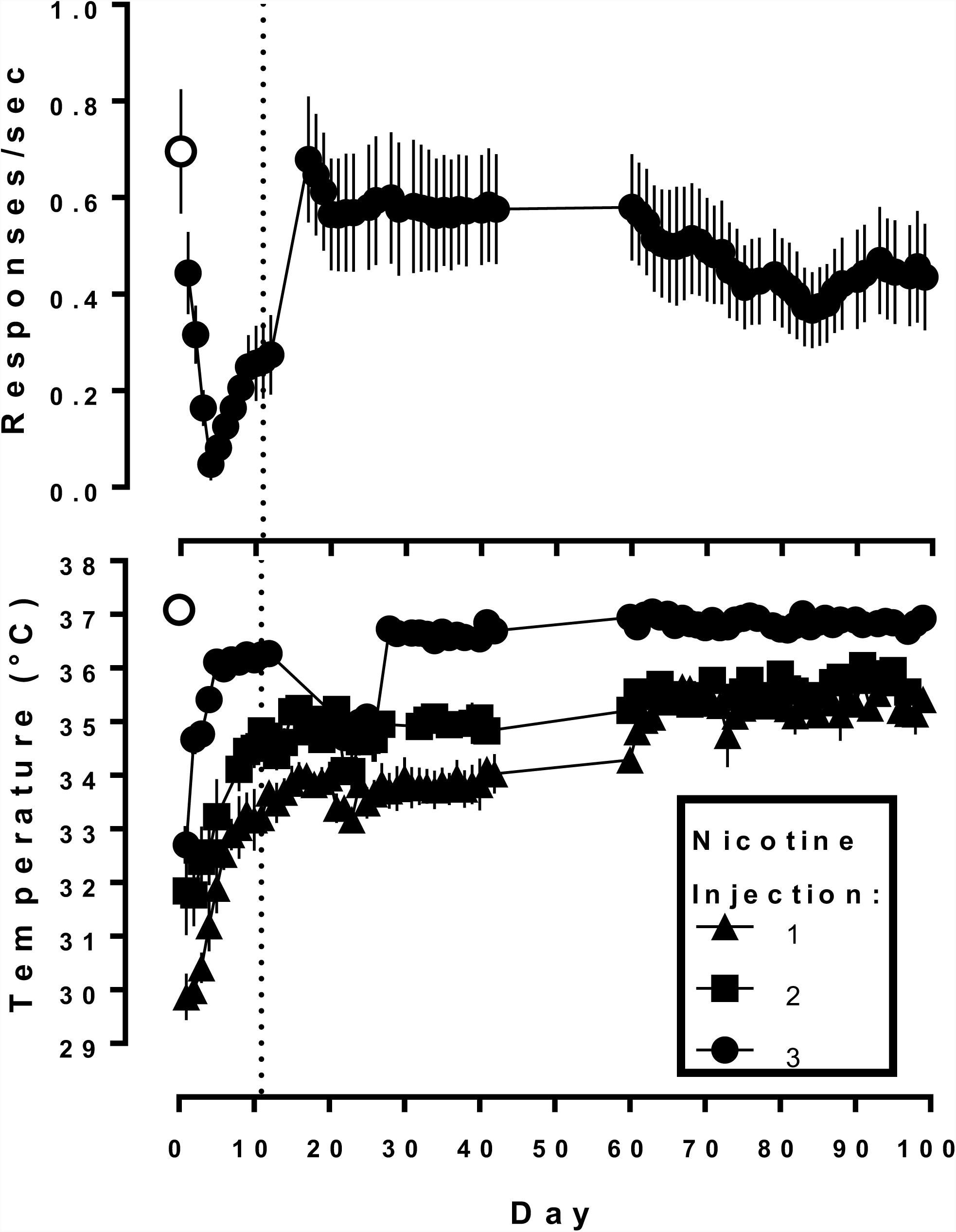
Response rate (top panel) and rectal temperature (bottom panel) during daily nicotine exposure with 3 doses of 1.78 mg/kg nicotine administered 90 min apart. For days 1-10, the third dose was administered at the beginning of operant conditioning sessions. From session 11 onward (vertical dashed line), operant conditioning sessions began 1 h after the second nicotine dose. Temperature was measured 30 min after each dose throughout the experiment. Only the effects of 1.78 mg/kg nicotine are shown, i.e. data from administration of other doses and drugs are omitted. Ordinate response rate expressed responses/sec (top panel), and rectal temperature expressed as °C (bottom panel). Abscissa: days of daily nicotine exposure.

**Figure 4:**
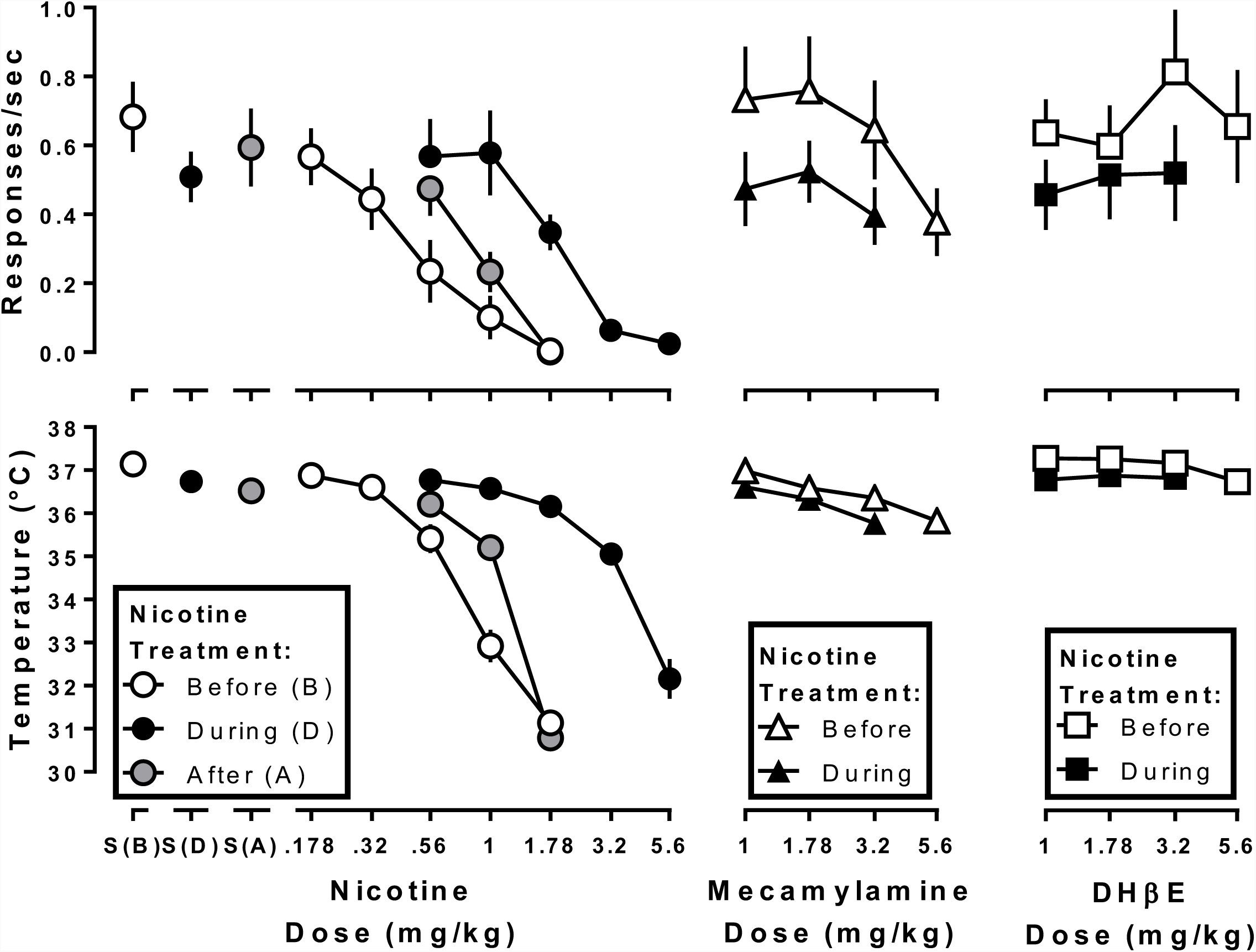
Rate of responding (top panels) and rectal temperature (bottom panels) following nicotine, mecamylamine, and DHβE before (open symbols) and during (filled symbols) daily nicotine exposure. The gray circles are the effects of nicotine after discontinuation of daily nicotine exposure. Ordinate: response rate expressed in responses/s (top panels), and rectal temperature expressed as °C (bottom panels). Abscissa: drug dose in mg/kg body weight.

Mecamylamine, when studied alone and prior to daily nicotine exposure, significantly decreased response rate to 0.38 responses per s at a dose of 5.6 mg/kg (F_4,24_=8.7, p=0.0023) (Figure 4, open triangles). Mecamylamine (3.2 and 5.6 mg/kg) significantly decreased rectal temperature as compared with saline control (Figure 4, open circle above S(B)) (F_4,24_=25.5, p=0.0002), with the largest decrease to 35.8°C. During daily nicotine exposure, mecamylamine no longer produced significant decreases in response rate relative to the corresponding saline control (F_3,18_=0.27, p=0.27) (Figure 4, filled circle above S(D) and filled triangles), but did produce hypothermic effects at doses of 1.78 and 3.2 mg/kg as compared with the daily nicotine treatment saline control (Figure 4, filled circle above S(D)) (F_3,18_=41.4, p<0.0001). When normalized to their respective saline controls determined before and during nicotine treatment, the mecamylamine dose-effect functions for decreasing response rate and rectal temperature were not significantly different (F_2,32_=0.21, p=0.21 and F_2,20_=0.64, p=0.54, respectively).

DHβE, when studied prior to daily nicotine treatment, did not significantly decrease response rate (F_4,24_=1.2, p=0.33), but did produce a small yet significant decrease in rectal temperature at 5.6 mg/kg (F_4,24_=5.7, p=0.025) (Figure 4, open squares). During daily nicotine exposure, DHβE (up to 3.2 mg/kg) did not significantly alter response rate (F_3,18_=0.30, p=0.69) or rectal temperature (F_3,18_=0.65, p=0.49) (Figure 4, filled squares). When normalized to the respective saline controls, the dose-effect functions of DHβE to decrease rectal temperature before versus during daily nicotine exposure were not significantly different from each other (F_2,24_=1.8, p=0.19).

During daily nicotine exposure, the dose-effect function for nicotine to decrease response rate in the presence of 3.2 mg/kg mecamylamine was significantly different from the nicotine control dose-effect function (F_1,6_=13.6, p=0.01) (Figure 5, top left). A Sidak’s post-hoc test demonstrated that response rate at only the 1 mg/kg dose of nicotine differed significantly in the presence versus the absence of 3.2 mg/kg mecamylamine (p<0.05). A Dunnett’s multiple comparison test revealed that response rate after 1 mg/kg nicotine in combination with 3.2 mg/kg mecamylamine did not significantly differ from that after mecamylamine alone (p>0.05). In contrast to response rate, 3.2 mg/kg mecamylamine significantly antagonized the hypothermic effects of nicotine (F_2,34_=9.2, p=0.0006). The dose-effect functions for nicotine alone and nicotine in combination with 3.2 mg/kg DHβE were significantly different from each other for rate-decreasing (F_1,6_=16, p=0.008) and hypothermic effects (F_2,38_=13, p<0.0001) (Figure 5, right). A Sidak’s post-hoc test indicated that the rate-decreasing effects of 0.56 and 1 mg/kg nicotine were significantly different following pretreatment with 3.2 mg/kg DHβE as compared to pretreatment with saline (p<0.05). However, a Dunnett’s post hoc test demonstrated that the rate-decreasing effects of 0.56 and 1 mg/kg nicotine in the presence of 3.2 mg/kg DHβE were not significantly different from response rate after DHβE alone (p>0.05).

**Figure 5:**
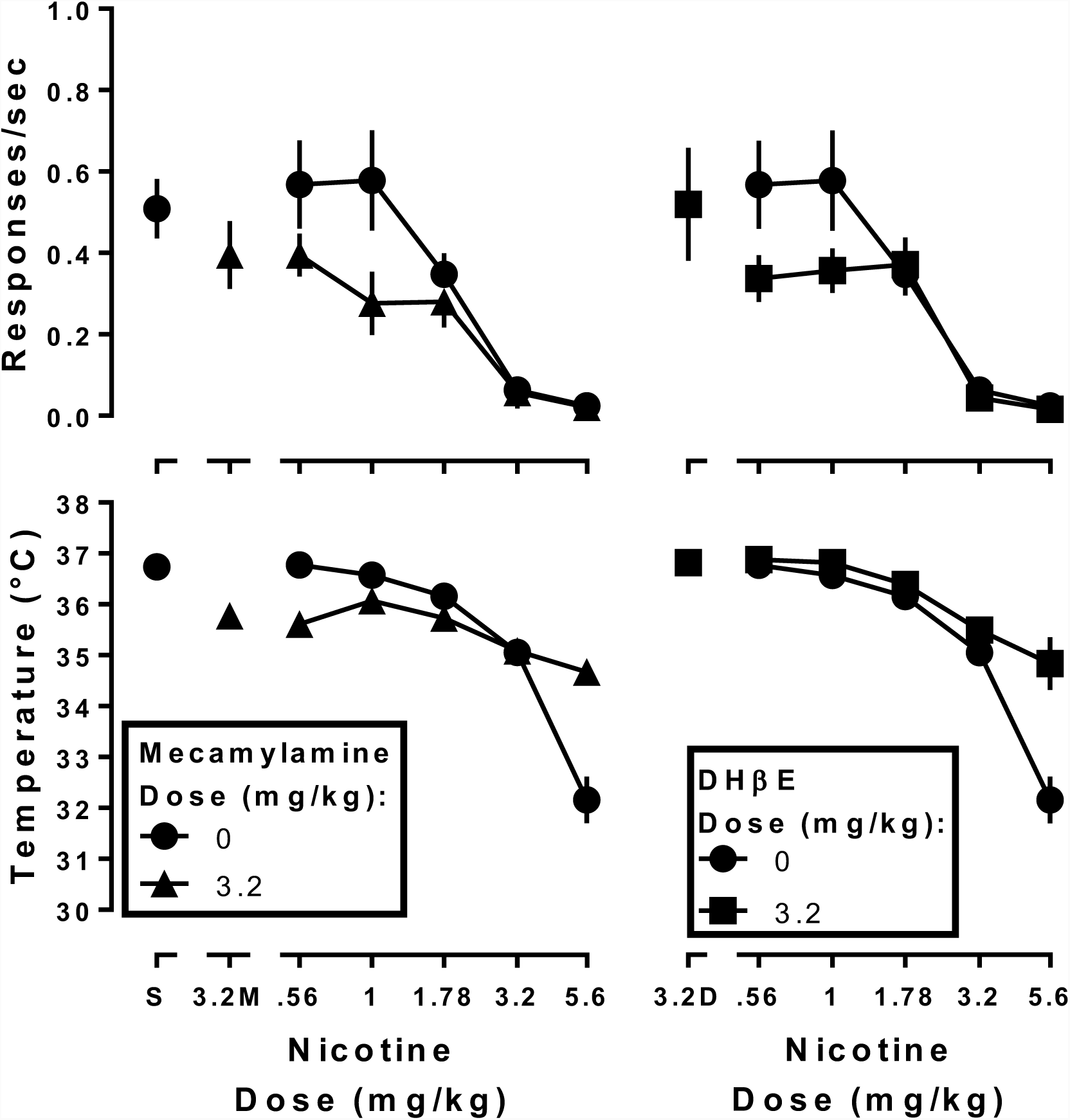
Response rate (top panels) and rectal temperature (bottom panels) following nicotine alone (filled circles) or in combination with 3.2 mg/kg mecamylamine (left panels; filled triangles) or DHβE (right panels; filled squares) during daily nicotine exposure. Ordinate: response rate expressed in responses/s (top panel), and rectal temperature expressed as °C (bottom panel). Abscissa: drug dose in mg/kg body weight.

Following discontinuation of daily nicotine treatment, the effects of 1 mg/kg nicotine were determined every 8 days. The effects of 1 mg/kg nicotine on day 32 were not significantly different from the effects of nicotine on days 16 and 24 for both rate-decreasing (F_2,12_=0.72, p=0.45) and hypothermic effects (F_2,12_=0.59, p=0.48). The nicotine dose-effect function was re-determined beginning on day 33 after discontinuation from daily nicotine treatment. The dose-effect function for nicotine to produce rate-decreasing (F_2,12_=58.7, p<0.0001) and hypothermic effects (F_2,12_=39.5, p=0.0007) significantly differed among the three conditions (i.e., before, during, and after discontinuation of nicotine treatment). The ED_50_ values for nicotine to produce rate-decreasing and hypothermic effects following discontinuation of daily nicotine treatment were 0.74 (0.57-0.96) and 1.1 (0.93-1.2) mg/kg, respectively (Figure 4, left panels, circles). Nicotine was 2.4 (1.6-3.4) and 3.0 (2.4-3.6) fold more potent to produce rate-decreasing and hypothermic effects, respectively, after discontinuation of daily nicotine, than during daily nicotine exposure. However, nicotine was 1.7 (1.2-2.4) and 1.6 (1.3-1.8) fold more potent to produce rate-deceasing and hypothermic effects, respectively, prior to daily nicotine exposure than after daily nicotine exposure.

After discontinuation of daily nicotine exposure, mecamylamine (1 mg/kg) significantly antagonized the rate-decreasing (F_2,32_=8.0, p=0.0015) and hypothermic effects (F_2,34_=54.3, p<0.0001) of nicotine (Figure 6, left). Mecamylamine shifted the nicotine dose-effect function for rate-decreasing effects 1.9 (1.5-2.5) fold to the right. Because of a significant difference in the slopes of the dose-effect functions for producing hypothermia (F_1,34_=19.3, p=0.0001), a potency ratio for nicotine alone versus nicotine in combination with 1 mg/kg mecamylamine was not determined. DHβE antagonized the rate-decreasing (F_2,34_=7.4, p=0.0021) and hypothermic effects of nicotine (F_2,32_=32.4, p<0.0001) following discontinuation of daily nicotine exposure (Figure 6, right). DHβE shifted the nicotine dose-effect functions 2.1 (1.5-2.8) and 2.6 (2.1-3.1) fold to the right for rate-decreasing and hypothermic effects, respectively. DHβE significantly differed in its antagonism of nicotine before daily nicotine exposure as compared with after discontinuation of daily nicotine exposure for both rate-decreasing (F_2,12_=20.1, p=0.001) and hypothermic effects (F_2,12_=46.8, p<0.0001), with post-hoc analysis indicating that antagonism of nicotine by DHβE was significantly less after discontinuation of daily nicotine exposure than before daily nicotine exposure.

**Figure 6:**
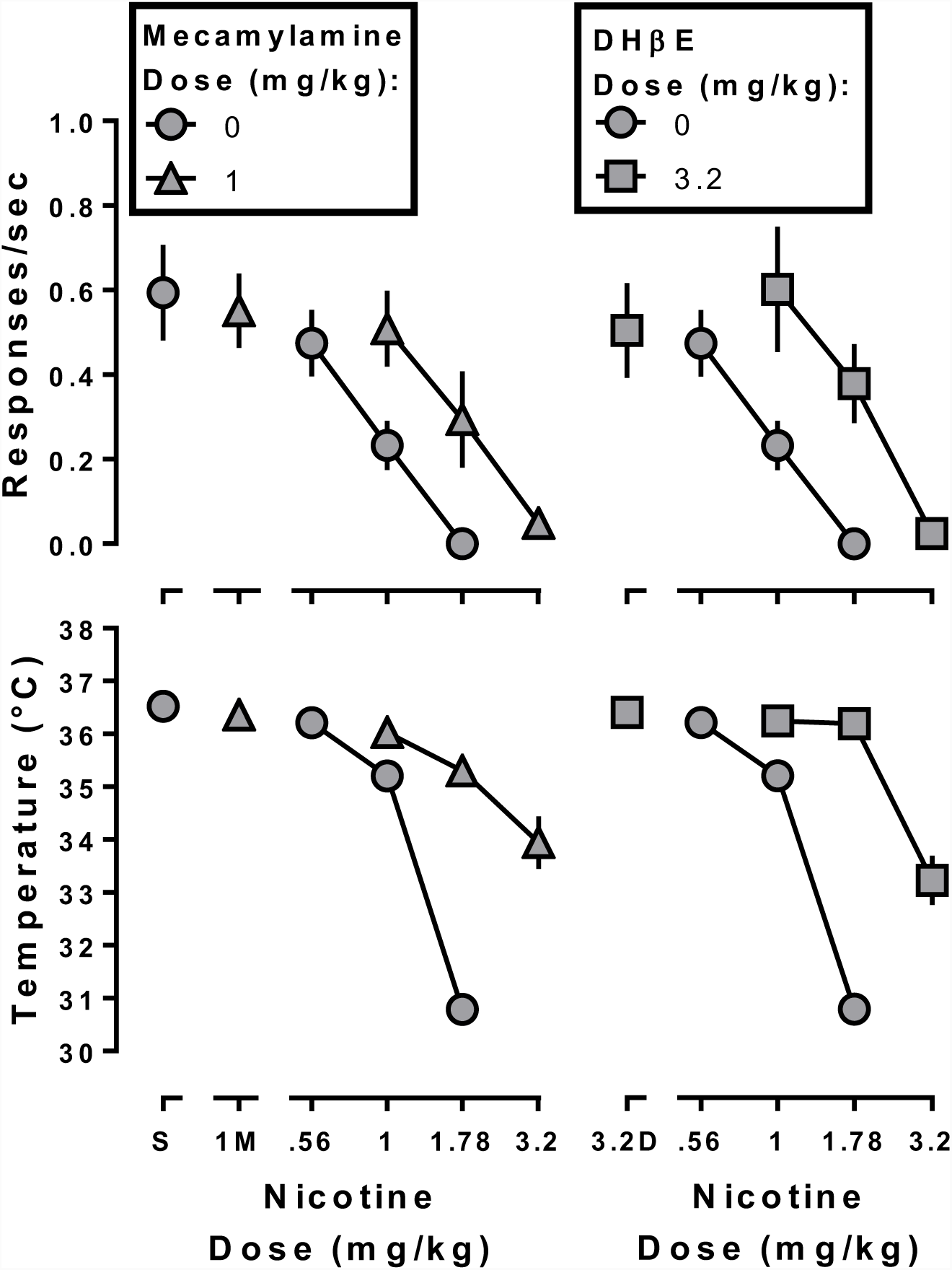
Response rate (top panels) and rectal temperature (bottom panels) following nicotine alone (gray circles) or in combination with 3.2 mg/kg mecamylamine (left panels; gray triangles) or DHβE (right panels; gray squares) after discontinuation of daily nicotine exposure. Ordinate: response rate expressed in responses/s (top panel), and rectal temperature expressed as °C (bottom panel). Abscissa: drug dose in mg/kg body weight. The control nicotine dose-effect functions shown in the left and right panels are the same.

## Discussion

Nicotine dose-dependently decreased responding under a fixed ratio schedule of liquid food delivery and decreased rectal temperature before, during, and after daily nicotine exposure. Significant tolerance developed to these effects of nicotine during daily nicotine exposure as evidenced by a rightward shift in the nicotine dose-effect functions. Before daily nicotine exposure, the effects of nicotine were antagonized by DHβE. The capacity of mecamylamine to antagonize nicotine prior to daily nicotine exposure was not studied because mecamylamine has been repeatedly demonstrated to antagonize the rate-decreasing and hypothermic effects of nicotine under conditions identical to or similar with those here (Cunningham and McMahon, 2011; Rodriguez et al., 2014; de Moura and McMahon, 2016). During daily nicotine exposure, mecamylamine and DHβE no longer antagonized the rate-decreasing effects of nicotine; however, the hypothermic effects of nicotine were still antagonized. Following discontinuation of daily nicotine exposure, both the rate-decreasing and hypothermic effects of nicotine were antagonized by DHβE and mecamylamine. However, the degree to which DHβE shifted the nicotine dose-effect functions rightward was significantly greater before daily nicotine exposure than after daily nicotine exposure. These results imply the following: the rate-decreasing and hypothermic effects of nicotine are mediated by a different receptor types or mechanisms, the rate-decreasing effects of nicotine in nicotine-tolerant animals may be mediated by non-nAChRs, and the antagonist actions of nAChR drugs are compromised in nicotine-tolerant and perhaps nicotine-dependent individuals (i.e. cigarette smokers).

As with many other drug classes (e.g., opioids), disruption of operant behavior has been used to examine precipitated withdrawal in dependent animals (Schulteis et al., 1994; Vann et al., 2006). Based on previous research demonstrating that 6 mg/kg/day of nicotine would not produce observable signs of withdrawal in C57BL/6J mice (Damaj et al., 2003), we predicted that our protocol would not increase sensitivity to the rate-decreasing effects of mecamylamine and DHβE. As expected, there was no increased sensitivity to the rate-decreasing and hypothermic effects of DHβE and mecamylamine, suggesting that the current regimen does not result in robust physical dependence.

Antagonism of nicotine by DHβE before and after chronic nicotine are consistent with the results of previous studies (de Moura and McMahon, 2016) and suggest that the rate-decreasing and hypothermic effects of nicotine are mediated by β2-containing nAChRs. Failure of DHβE and mecamylamine to antagonize the rate-decreasing effects of nicotine during daily nicotine exposure suggests that in nicotine-tolerant animals, the rate-decreasing effects of nicotine are produced by actions at non-nAChRs. In contrast, the hypothermic effects of nicotine continue to be mediated by β2 nAChRs as evidenced by antagonism of nicotine by both mecamylamine and DHβE. Because daily nicotine exposure decreases functional nAChRs (Quick and Lester, 2002; Giniatullin et al., 2005), lack of antagonism of the rate-decreasing effects of nicotine by mecamylamine and DHβE suggests that nicotine may act at other receptor types (e.g., non-nAChR) to disrupt behavior. Differential antagonism of the rate-decreasing and hypothermic effects of nicotine suggests that different receptors mediate these effects.

The current results are consistent with previous research demonstrating that multiple nAChRs mediate the *in vivo* effects of nicotine (Stolerman et al., 1997; de Moura and McMahon, 2016). For instance, the nAChR agonist cytisine will only partially substitute for nicotine in rats trained to discriminate 0.32 mg/kg nicotine; however, when the training dose of nicotine is increased to 1.78 mg/kg, cytisine will fully substitute for nicotine (Jutkiewicz et al., 2011). This result suggests that nAChR subtypes differentially mediate the discriminative stimulus effects of nicotine as a function of training dose, where low doses of nicotine may activate only a subset of the nAChR subtypes that are activated at higher doses. Furthermore, Stolerman et al. (1997) demonstrated that DHβE differentially antagonizes various behavioral effects of nicotine. Of particular note, DHβE antagonized the discriminative stimulus effects of nicotine, but not its rate-decreasing effects (Stolerman et al., 1997). From these results, the authors concluded that nAChR subtypes differentially mediate the behavioral effects of nicotine. The failure of DHβE to antagonize the rate-decreasing effects of nicotine in Stolerman et al. (1997) might suggest that tolerance had developed to the rate-decreasing effects of nicotine over the course of discrimination training and testing, thereby decreasing the ability of DHβE to antagonize rate-decreasing effects.

That nicotine was more potent to decrease response rate and rectal temperature before as compared with approximately 5 weeks after discontinuation of daily nicotine treatment is consistent with previously published reports (Rosecrans et al., 1989). The loss of potency to the effects of nicotine on schedule-controlled responding even after several weeks of discontinued daily nicotine treatment could reflect behavioral tolerance; however, because tolerance to nicotine under these treatment conditions is greater than cross-tolerance to cytisine and cocaine, as reported previously (de Moura and McMahon, 2016), pharmacodynamic changes at the level of nAChRs (i.e., desensitization) appear to play an important role. Moreover, the persistent loss of potency to a physiological effect of nicotine (i.e., hypothermia) suggests that behavioral tolerance is insufficient to explain the results obtained with schedule-controlled responding. The persistent loss of nicotine’s potency as a consequence of daily nicotine treatment, coupled with the decreased effectiveness of DHβE as an antagonist, suggests that neuroadaptations in nAChR signaling persist long after chronic treatment. This result is consistent with studies that demonstrated changes in receptor expression levels following various amounts of exposure to nicotine (Olale et al., 1997; Fenster et al., 1999; Buisson and Bertrand, 2001; Nashmi et al., 2007).

While there is interest in using nAChR antagonists to treat tobacco use in a manner similar to that underlying the use of the µ-opioid antagonist naltrexone as a treatment for opioid use disorder, these results suggest that antagonists may be limited in their capacity to function as smoking cessation aids. It is unclear why mecamylamine and DHβE are no longer able to antagonize the rate-decreasing effects of nicotine after daily nicotine exposure. It is possible that these effects of nicotine are mediated by non-nAChRs in tolerant mice, or that repeated nicotine exposure changes how mecamylamine and DHβE interact with the nAChR. However, these results suggest that multiple nAChRs mediate the various in vivo effects of nicotine, which is consistent with previously published reports (Stolerman et al., 1997; de Moura and McMahon, 2016). Future research should attempt to identify which receptors mediate the effects of nicotine during repeated nicotine exposure. Identification of these receptors may provide a blueprint in the development of more effective tobacco cessation pharmacotherapies.

## Acknowledgements

The authors would like to thank Morgan Cocke and Julia Threadgill for their technical assistance, and Dr. Brett Ginsburg for suggestions for data analyses. This study was funded by USPHS DA25267.

